# Double maternal effect: duplicated nucleoplasmin 2 genes, *npm2a* and *npm2b*, are shared by fish and tetrapods, and have distinct and essential roles in early embryogenesis

**DOI:** 10.1101/104760

**Authors:** Caroline T. Cheung, Jérémy Pasquier, Aurélien Bouleau, Thao-Vi Nguyen, Franck Chesnel, Yann Guiguen, Julien Bobe

**Affiliations:** INRA LPGP UR1037, Campus de Beaulieu, 35042 Rennes, FRANCE.; CNRS/ UMR6290, Université de Rennes 1, 35000 Rennes, FRANCE.

## Abstract

*Nucleoplasmin 2* (*npm2*) is an essential maternal-effect gene that mediates early embryonic events through its function as a histone chaperone that remodels chromatin. Here we report the existence of two *npm2* (*npm2a* and *npm2b*) genes in zebrafish. We examined the evolution of *npm2a* and *npm2b* in a variety of vertebrates, their potential phylogenetic relationships, and their biological functions using knockout models via the CRISPR/cas9 system. We demonstrated that the two *npm2* duplicates exist in a wide range of vertebrates, including sharks, ray-finned fish, amphibians, and sauropsids, while *npm2a* was lost in Coelacanth and mammals, as well as some specific teleost lineages. Using phylogeny and synteny analyses, we traced their origins to the early stages of vertebrate evolution. Our findings suggested that *npm2a* and *npm2b* resulted from an ancient local gene duplication, and their functions diverged although key protein domains were conserved. We then investigated their functions by examining their tissue distribution in a wide variety of species and found that they shared ovarian-specific expression, a key feature of maternal-effect genes. We also showed that both *npm2a* and *npm2b* are maternally-inherited transcripts in vertebrates. Moreover, we used zebrafish knockouts to demonstrate that *npm2a* and *npm2b* play essential, but distinct, roles in early embryogenesis. *npm2a* functions very early during embryogenesis, at or immediately after fertilization, while *npm2b* is involved in processes leading up to or during zygotic genome activation. These novel findings will broaden our knowledge on the evolutionary diversity of maternal-effect genes and underlying mechanisms that contribute to vertebrate reproductive success.

**Author Summary:** The protein and transcript of the *npm2* gene have been previously demonstrated as maternal contributions to embryos of several vertebrates. Recently, two *npm2* genes, denoted here as *npm2a* and *npm2b*, were discovered in zebrafish. This study was conducted to explore the evolutionary origin and changes that occurred that culminated in their current functions. We found that an ancient local duplication of the ancestral *npm2* gene created the current two forms, and while most vertebrates retained both genes, notably, mammals and certain species of fish lost *npm2a* and, albeit rarely, both *npm2a* and *npm2b*. Our functional analyses showed that *npm2a* and *npm2b* have diverse but essential functions during embryogenesis, as *npm2a* mutants failed to undergo development at the earliest stage while *npm2b* mutants developed, although abnormally, until the zygotic genome activation stage after which their development was arrested followed subsequently by death. Our study is the first to clearly demonstrate the evolution, diversification, and functional analyses of the *npm2* genes, which are essential maternal factors that are required for proper embryonic development and survival.

## Introduction

In animals and plants, early embryonic development relies strictly on maternal products until maternal-to-zygotic transition (MZT) during which zygotic genome activation (ZGA) occurs [1]. Maternal-effect genes are those that are transcribed from the maternal genome and whose products, which include transcripts, proteins, and other biomolecules, are deposited into the oocytes during their production in order to coordinate embryonic development before ZGA [2]. MZT is a key step that is needed firstly for clearance of maternal components, and secondly to activate zygotic gene expression and to allow subsequent embryonic development. Among the maternally-inherited transcripts that play important roles during early development, some genes were demonstrated to regulate zygotic program activation such as *nanog*, *pou5f1*, and *soxb1* in zebrafish (*Danio rerio*) [3]. Henceforth, all gene and protein nomenclature will be based on that of zebrafish regardless of species for simplification purposes. Those factors were shown to participate in the regulation of “first wave” zygotic genes and among them, mir-430, a conserved microRNA that has been shown to be involved in clearance of maternal mRNAs in zebrafish [4][5], as well as mir-427, its orthologue in Xenopus [6]. microRNAs may also have maternal effects by functioning in a similar manner to protein-coding genes, and recent revelations showed that there is a subset of microRNAs that are predominantly expressed in the fish ovary and may function in oogenesis and early embryogenesis [7]. Another gene, *nucleoplasmin 2* (*npm2*), belongs to the family of nucleoplasmins/nucleophosmins that is maternally-inherited at both protein and mRNA levels, whereby both play important roles in early development [8]. Historically, this protein was identified and defined as a nuclear chaperone in Xenopus [9,10]. While the protein has been shown to be the most abundant nuclear protein in the Xenopus oocyte [11] and to play a crucial role at fertilization due to its role in sperm chromatin decondensation [12,13], its maternally-inherited mRNA has been recently demonstrated to be translated as a *de novo* synthesized protein that could play a crucial role during ZGA in zebrafish [8]. Further, *npm2* is one of the first identified maternal-effect genes in mouse whereby its deficiency results in developmental defects and eventual embryonic mortality [14].

The *npm2* gene belongs to the *npm* gene family that encompasses four members, *npm1, npm2, npm3*, *and npm4*. The diversity of the Npm family has been shown to result from the two rounds of whole genome duplication (WGD) that occurred in early vertebrates (vertebrate genome duplication 1 and 2, or VGD1 & VGD2, respectively) [15,16]. Former evolutionary studies clearly provided a phylogenic model of this family; VGD1 produced two genes, *npm3/2* and *npm4/1*, from an ancestral *npm* gene and the following WGD, VGD2, further created the current four *npm* types with subsequent loss of *npm4* from mammals, but retained in most fish species [17–20]. Recently, two *npm2* genes were automatically annotated in the zebrafish genome, i.e. *npm2a* (ENSDARG00000076391) and *npm2b* (previously known as *npm2*, ENSDARG00000053963). As the teleost ancestor experienced an extra WGD event (TGD, or teleost-specific genome duplication) [21], doubling of genes and other types of genomic rearrangement may be present in teleost species compared with other vertebrates. Moreover, a fourth round of duplication occurred more recently in salmonids (salmonid-specific genome duplication or SaGD) [22,23], leading to further possible doubling of genes and other genomic rearrangements. As multiple evolutionary events (impact of TGD, local duplication, teleost-specific duplication, etc.) could have led to the *npm2a*/*npm2b* diversity in zebrafish, which in turn may have significant impact on vertebrate reproduction, we investigated the evolution of *npm2a* and *npm2b* in a wide range of vertebrate species, their potential phylogenetic relationship, as well as their biological functions using transgenic zebrafish models created by the CRISPR/cas9 system in order to broaden our knowledge on the evolutionary diversity of maternal-effect genes and the underlying mechanisms that contribute to reproductive success in vertebrates.

## Results and Discussion

As previously shown, *npm1*, *npm2*, *npm3* and *npm4* genes are thought to have originated from the first two rounds of WGD, VGD1 and VGD2, which occurred early on in vertebrate evolution [17–20]. However, two *npm2* genes have been automatically annotated in the zebrafish genome, *i.e. npm2a* and *npm2b*. In order to verify if these two *npm2* genes are paralogous to each other and to determine their origination, we used a Blast search approach in various public databases to retrieve a multitude of sequences that could be related to *npm2* genes. All retrieved sequences are compiled in Supplemental Table S1.

### Diversity of npm2a and npm2b in vertebrates

#### Phylogenetic analysis

In order to verify that the retrieved protein sequences (Supplemental Table S1) were homologous to zebrafish Npm2a and Npm2b, a phylogenetic analysis on Npm2 was performed. Based on the alignment of 76 vertebrate Npm2-related sequences, and using vertebrate Npm1 and Npm3 amino acid sequences as out-groups, a phylogenetic tree was generated (Fig. 1). As shown in Fig. 1, vertebrate Npm2-related sequences clustered into two clades, Npm2a and Npm2b, which were supported by significant bootstrap values, 84.2% and 78.2%, respectively.

**Figure 1:**
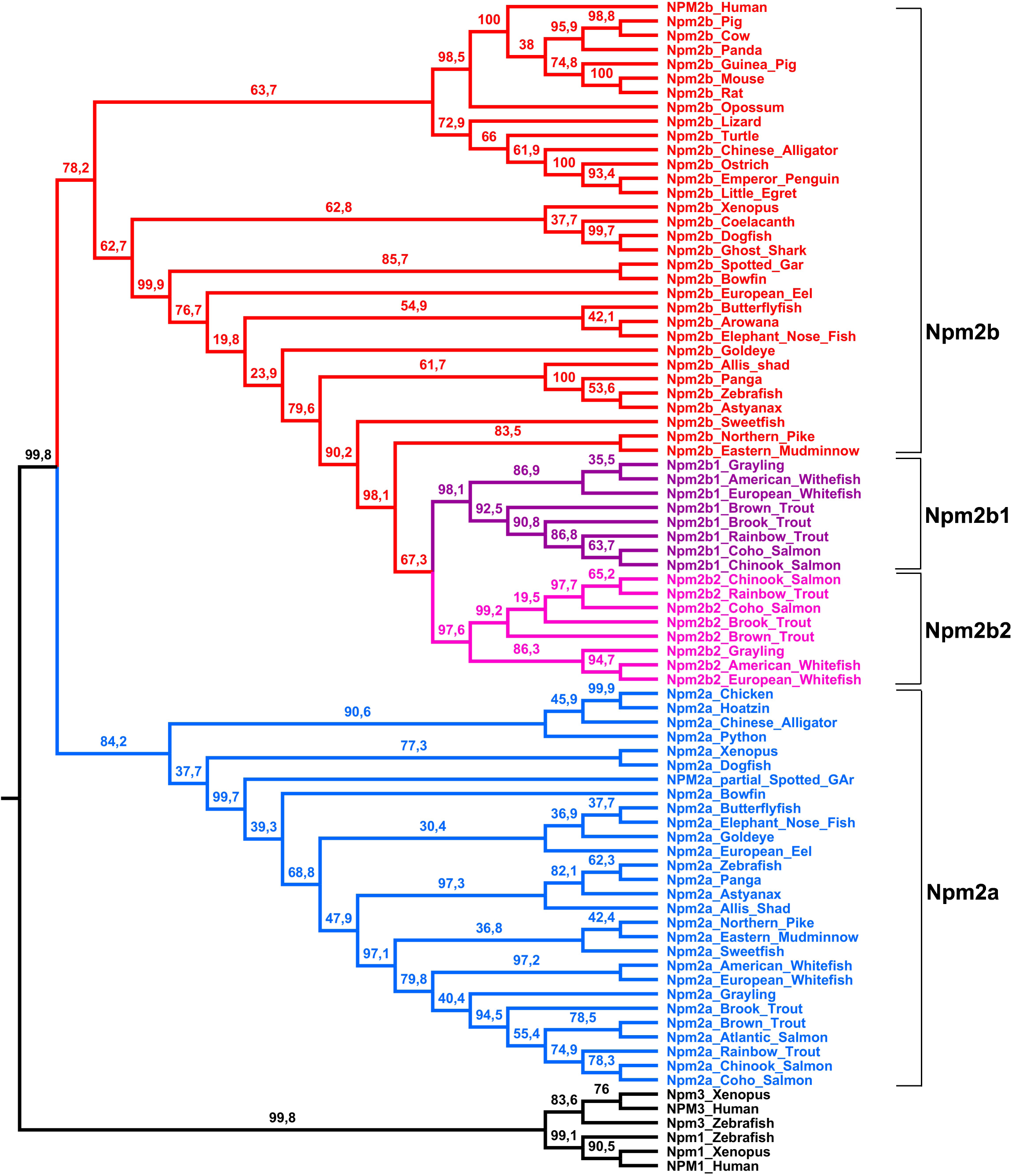
Consensus phylogenetic tree of Npm2 proteins. This phylogenetic tree was constructed based on the amino acid sequences of Npm2 proteins (for the references of each sequence see Supplemental Table S1) using the Neighbour Joining method with 1,000 bootstrap replicates. The number shown at each branch node indicates the bootstrap value (%). The tree was rooted using Npm3 and Npm1 sequences. The Npm2a sequences are in blue, the Npm2b sequences are in red, the salmonid Npm2b1 sequences are in purple, and the salmonid Npm2b2 sequences are in pink.

There were 28 sequences that clustered in the Npm2a clade, which encompassed sequences belonging to species from various vertebrate groups, including chondrichthyans (such as dogfish Npm2a), ray-finned fish, and sarcopterygians (lobe-finned fish and tetrapods). The sequences belonging to ray-finned fish included gar and bowfin Npm2a, as well as teleost sequences such as zebrafish, northern pike, and rainbow trout Npm2a. In contrast, no Npm2a sequence was identified in neoteleostei (medaka, European perch, and Atlantic cod). The sequences belonging to sarcopterygians included amphibian (Xenopus) Npm2a, as well as sauropsid sequences such as Chinese alligator, chicken, penguin, and ibis Npm2a, although no Npm2a sequence could be identified in mammals. These results provided the first evidence for the presence of orthologs to zebrafish Npm2a across evolutionary divergent vertebrate groups (i.e. sauropsids, amphibians, and fish).

The Npm2b clade included 48 of the retrieved sequences. Like the Npm2a clade, the Npm2b clade also encompassed sequences belonging to species from all of the main vertebrate groups, including chondrichthyans (such as dogfish Npm2b), ray-finned fish, and sarcopterygians. Ray-finned fish sequences included gar and bowfin Npm2b in addition to teleost sequences such as zebrafish and northern pike Npm2b. Our analysis also demonstrated the existence of two paralogous Npm2b proteins (*i.e.* Npm2b1 and Npm2b2) in all of the investigated salmonid species (rainbow trout, brown trout, and brook trout). In contrast, no Npm2b sequence could be identified in any neoteleostean species, including medaka, cod, perch, fugu, tetraodon, and stickleback. The sequences belonging to sarcopterygians included amphibian sequences such as Xenopus Npm2b, sauropsid sequences such as Chinese alligator, ostrich, penguin and chicken Npm2b, as well as mammalian sequences such as human and mouse Npm2. These results demonstrated, for the first time, that proteins previously reported as Npm2 (Xenopus, cattle, mouse, human, zebrafish) are orthologous to Npm2b in all investigated vertebrate species, and should therefore now be referred to as Npm2b. In addition, with the presence of Npm2b1 and Npm2b2 in all investigated salmonid species, we provided for the first time evidence of the existence of two Npm2b protein forms in vertebrate species.

The existence of the two Npm2 clades indicated that Npm2a and Npm2b are likely paralogous to each other. In addition, comparison of Npm2a and Npm2b amino acid sequences in the species that harbor both revealed that they share between 30.2% and 46.4% homology, depending on the species (Supplemental Table S2). The low sequence identity is consistent with an ancient duplication event that gave rise to *npm2a* and *npm2*b genes. However, neither the topology of the Npm2 phylogenetic tree nor the comparison between Npm2a and Npm2b could indicate the kind of duplication event that had occurred. In contrast, the high sequence identity shared by Npm2b1 and Npm2b2 in salmonids (between 75.1% and 86%) suggested a more recent duplication event. This observation in all of the investigated salmonid species is consistent with the hypothesis that *npm2b1* and *npm2b2* genes likely resulted from the SaGD.

#### Synteny analysis

In order to further understand the origin of the *npm2a* and *npm2b* genes in vertebrates, we performed a synteny analysis of their neighboring genes in representative vertebrate genomes. We focused our study on two mammals (human and mouse), two sauropsids (Chinese alligator and chicken), two amphibians (*Xenopus tropicalis* and *laevi*s L), one basal sarcopterygian (coelacanth), one basal actinopterygian (spotted gar), and two teleosts (zebrafish and tetraodon) (Fig. 2). Analysis was also performed on the *Xenopus laevi*s S subgenome, but since the results were very similar to that of *Xenopus laevi*s L subgenome, we only showed the latter’s data [24].

**Figure 2:**
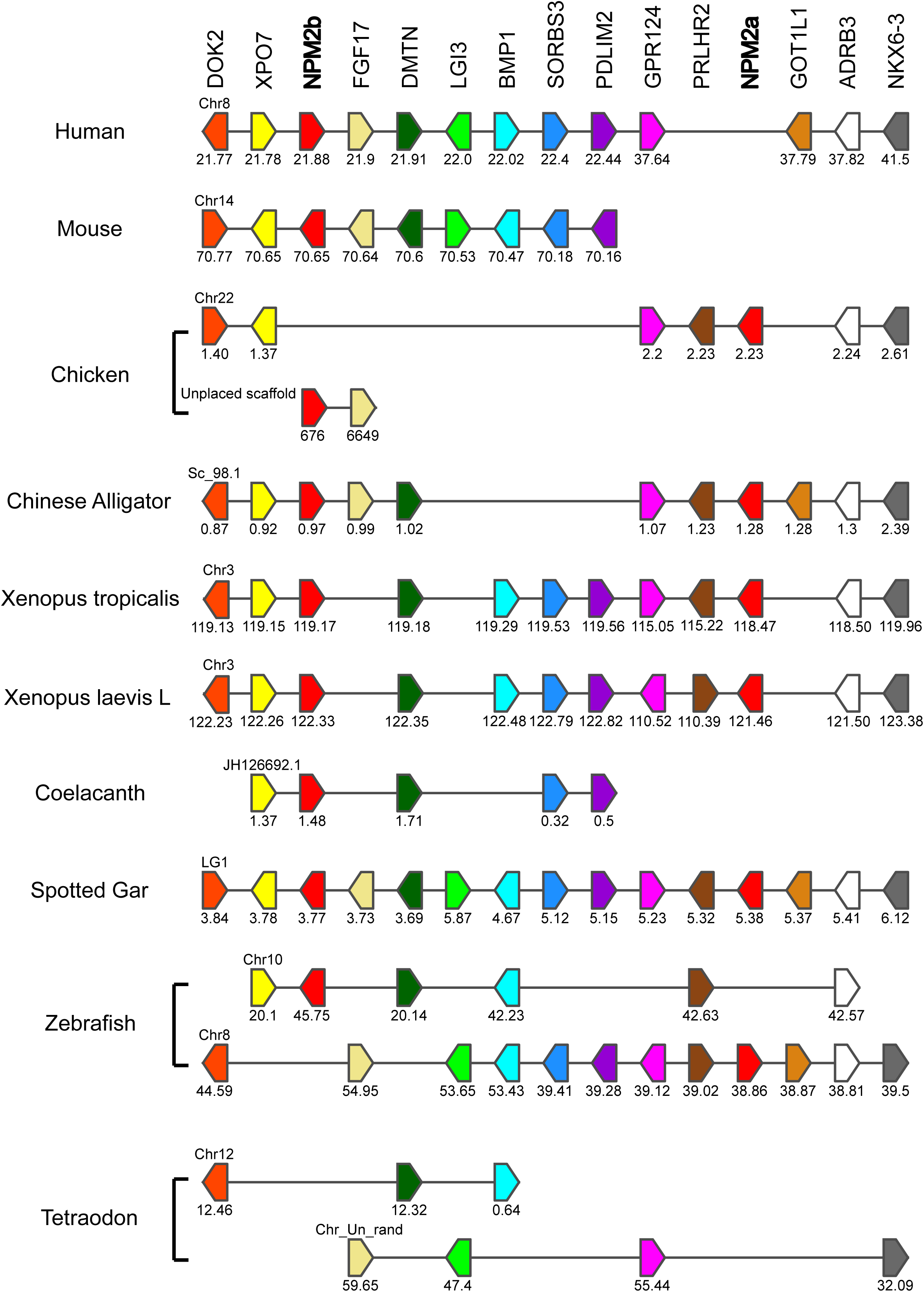
Conserved genomic synteny of *npm2* genes. Genomic synteny maps comparing the orthologs of *npm2a, npm2b*, and their neighbouring genes. *npm2* genes are named as *npm2a* and *npm2b* (formerly known as *npm2*). The other genes were named after their human orthologs according to the Human Genome Naming Consortium (HGNC). Orthologs of each gene are shown in the same color. The direction of arrows indicates the gene orientation, with the ID of the genomic segment indicated above and the position of the gene (in 10^-6^ base pairs) indicated below. The full gene names and detailed genomic locations are given in Supplemental Table S3.

The human, mouse, Chinese alligator, *Xenopus tropicalis* and *laevis*, coelacanth, spotted gar, and zebrafish *npm2b* genes are located in genomic regions containing common loci, including *dok2, xpo7, fgf17, dmtn, lgi3, bmp1, sorbs3, pdlim2, gpr124, prlhr, got1l1, adrb3,* and *nkx6-3*. Together with the phylogenetic analysis, this indicates that *npm2b* genes investigated here are orthologous. Synteny analysis revealed the absence of the *npm2b* gene in tetraodon although the above-mentioned neighbouring genes are present in its genome (Fig. 2 and Supplemental Table S3).

The Chinese alligator, chicken, *Xenopus tropicalis* and *laevis*, spotted gar, and zebrafish *npm2a* genes are located in genomic regions containing the same loci as *npm2b* conserved regions (Fig. 2). Indeed, Chinese alligator and spotted gar *npm2a* and *npm2b* genes are located in the vicinity of each other on scaffold 98.1 and the linkage group LG1, respectively. The presence of *npm2a* and *npm2b* genes in the same genomic region in representative species of sarcopterygians (Chinese alligator and *Xenopus tropicalis* and *laevis*) and actinopterygians (spotted gar) strongly suggested that *npm2a* and *npm2b* genes could have resulted from a unique local duplication of an ancestral *npm2* gene.

### Evolutionary history of npm2 genes in vertebrates

The presence/absence of *npm2a* and *npm2b* in the current vertebrate phyla and species is summarized in Fig.3, and we also propose an evolutionary scenario for the diversification of the *npm2* genes across vertebrate evolution.

**Figure 3:**
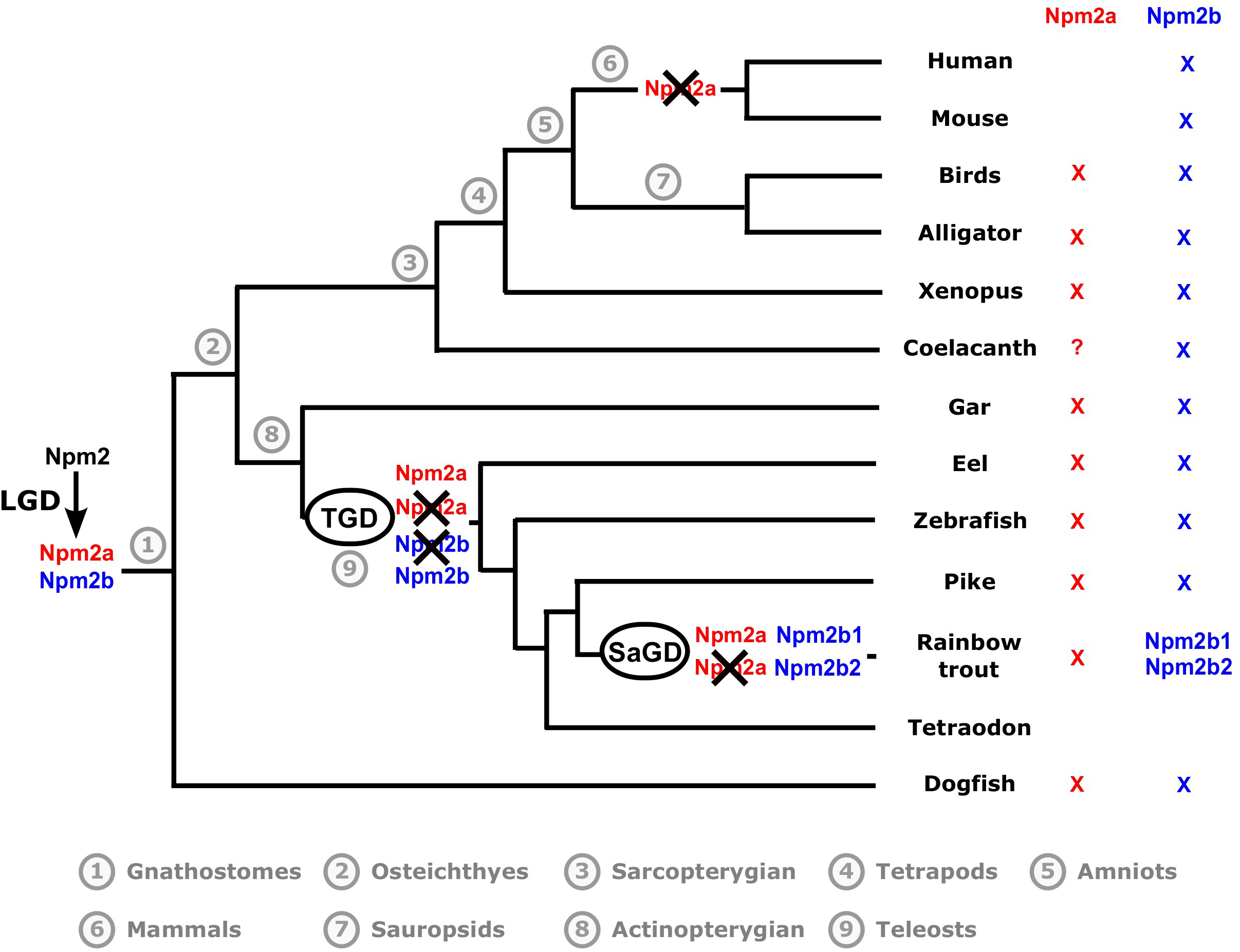
Current status and proposed evolutionary history of *npm2* genes among gnathostomes. The names of the current representative species of each phylum are given at the end of the final branches, together with a red and/or blue X to denote the *npm2* genes they possess (*npm2a*=red, *npm2b*=blue). The black X upon an *npm2* gene symbol indicates a gene loss. LGD: local gene duplication; TGD: teleost-specific whole genome duplication; SaGD: salmonid-specific whole genome duplication.

In this study, we demonstrated that *npm2a* and *npm2b* may be paralogous genes present in the different vertebrate groups, chondrichthyans (Fig. 1), sarcopterygians, and actinopterygians (Figs. 1 and 2), which strongly suggested that the *npm2* genes originated from a duplication event prior to the divergence of chondrichthyans and osteichthyans. Since the four *npm* family members (*npm1*, *npm2*, *npm3*, and *npm4*) are thought to be produced from the first two rounds of WGD (VGD1 & VGD2) that occurred early on in vertebrate evolution, we can thus hypothesize that the duplication event that generated *npm2a* and *npm2b* took place after VGD2, but before emergence of chondrichthyans and osteichthyans, between 450 and 500 million years ago (Mya) [25]. The findings from our synteny analysis demonstrated that in representative species of actinopterygians (spotted gar) and sarcopterygians (*Xenopus tropicalis* and *laevis* and Chinese alligator), *npm2a* and *npm2b* genes are at two distinct loci located on the same chromosomic region (Fig. 2). These results strongly suggested that *npm2a* and *npm2b* genes arose from local gene duplication rather than a whole genome (or chromosome) duplication event (Fig. 3).

The teleost ancestor experienced an extra WGD event (TGD) [21], but in all investigated teleosts, we observed a maximum of one *npm2a* ortholog and one *npm2b* ortholog (except in salmonids) (Fig. 1). In addition, *npm2a* and *npm2b* are located on two TGD ohnologous regions on chromosomes 8 and 10, respectively (Fig. 2), in zebrafish, which is consistent with the loss of one of the *npm2a* duplicates from one region and the loss of one *npm2b* duplicate from the corresponding ohnologous region. Thus, *npm2* diversity is most likely due to the early loss of one of the two *npm2a* and *npm2b* TGD ohnologs (Fig. 3). This observation in zebrafish strengthened the hypothesis that TGD did not impact the *npm2a* and *npm2b* diversity in teleosts. In addition, the lack of *npm2a* and *npm2b* in neoteleostei species, such as tetraodon, suggested that additional gene losses occurred early in the evolutionary history of this group (Fig. 2).

In salmonids, we identified two *npm2b* paralogs, i.e. *npm2b1* and *npm2b2*, in all investigated salmonid species (Fig. 1). Considering the high sequence identity shared by *npm2b1* and *npm2b2*, it is strongly hypothesized that these duplicates originated from SaGD. In contrast, we identified only one *npm2a* gene in all investigated salmonids suggesting that SaGD did not have any impact on the current salmonid *npm2a* diversity mostly due to an early loss of the SaGD ohnolog of this gene (Fig. 3).

In addition to the early gene losses after WGD, various other independent and phylum-specific gene losses may have contributed to shape the current diversity of *npm2a* and *npm2b* in vertebrates. In fact, although both *npm2a* and *npm2b* have been globally conserved in sarcopterygians and actinopterygians, some phyla in each group lack at least one of the genes. In sarcopterygians, *npm2a* is conserved in amphibians such as *Xenopus tropicalis* and *laevis*, and in sauropsids such as Chinese alligator and chicken (Figs. 1-3). In contrast, we did not find any *npm2a* genes in the Coelacanth genome (Figs. 1-3), which could be due to the lower assembly quality of the concerned genomic region (see Supplemental Table S3), and in the mammalian genomes including human and mouse (Figs. 1-3), whose absence may have been due to the loss of this gene in the common ancestor of mammals. On the other hand, *npm2b* is conserved in most investigated species. In actinopterygians, *npm2a* and *npm2b* are both conserved in all investigated species except neoteleostei in which neither *npm2a* nor *npm2b* is found. In contrast, one *npm2a* and two *npm2b*, i.e. *npm2b1* and *npm2b2*, are found in salmonids, representing to date the most important diversity of *npm2* genes in vertebrates (Figs. 1-3). Considering the essentialness of the *npm2* gene in embryonic development, its lack in some neoteleosteon species, such as Atlantic cod and medaka, raises the question on how evolution can cope with its loss. Further analyses of data from the Phylofish database [26] (data not shown) showed that *npm3* had the strongest homology to *npm2* and was predominantly expressed in the ovaries in medaka and cod which suggest that it could potentially compensate for *npm2* deficiency. In contrast, *npm3* does not show an ovarian-predominant expression in spotted gar and zebrafish, thus suggesting that its strong ovarian expression could be restricted to neoteleostei.

### Evolution of npm2a and npm2b coding sequences

To examine the evolutionary adaptation of the *npm2* proteins, estimation of the ratio of substitution rates (dN/dS) between the paralogous *npm2a* and *npm2b* CDS for all of the species harbouring multiple *npm2* paralogs was performed (Supplemental Table S4). For all investigated species, the dN/dS values were well below 1, indicating that whilst *npm2a* and *npm2b* genes diverged approximately 500 Mya, they have remained under strong purifying selection and evolutionary pressure, which may have tended to conserve their distinct protein structures and functions. Thus, in this study, we found that Npm2a and Npm2b may have diverged from an ancient duplication and currently share low sequence identity, suggesting that they could have thus evolved with different roles, which were likely conserved through strong negative selection.

## Expression domains of Npm2a and Npm2b in vertebrates

In order to investigate further the potential functions of Npm2a and Npm2b, we first explored the tissue distributions of both transcripts using two different approaches, qPCR and RNA-Seq, the latter of which was obtained from the Phylofish online database [26]. In zebrafish and *Xenopus tropicalis*, we observed that *npm2a* and *npm2b* were both predominantly expressed in the ovary, and to a lesser extent, the muscle, as well as in the zebrafish gills (Fig. 4A and 4B). We also demonstrated that *npm2a* and *npm2b* were predominantly expressed in the ovary of bowfin, elephantnose fish, panga, European eel, sweetfish, and northern pike (Fig. 4C to 4H). In the investigated salmonid species, brook trout and brown trout, *npm2a*, *npm2b1*, and *npm2b2* were also predominantly expressed in the ovary (Fig. 4I and 4J). In addition, *npm2* transcripts were expressed at a very low level in the testis of European eel, northern pike, and salmonid species (Fig. 4F, 4H, 4I, and 4J). In teleosts, *npm2a* mRNA levels globally tended to be lower than *npm2b* (or *npm2b1* and *npm2b2* for salmonids) (Fig. 4A and 4D-J). The clear ovarian-specific expression profiles of *npm2a* and *npm2b* transcripts were conserved in all investigated species, from teleosts to tetrapods. This is consistent with the *npm2b* mRNA profiles reported in the literature for mouse [14], cattle [27], *Xenopus tropicalis* [28], and zebrafish [8]. Despite the long passage of time since their divergence (approximately 500 Mya) [25] as demonstrated above, *npm2a* and *npm2b* appeared to have conserved their ovarian-specific expression profiles, which suggested that they both have roles in female reproduction and/or embryonic development.

**Figure 4:**
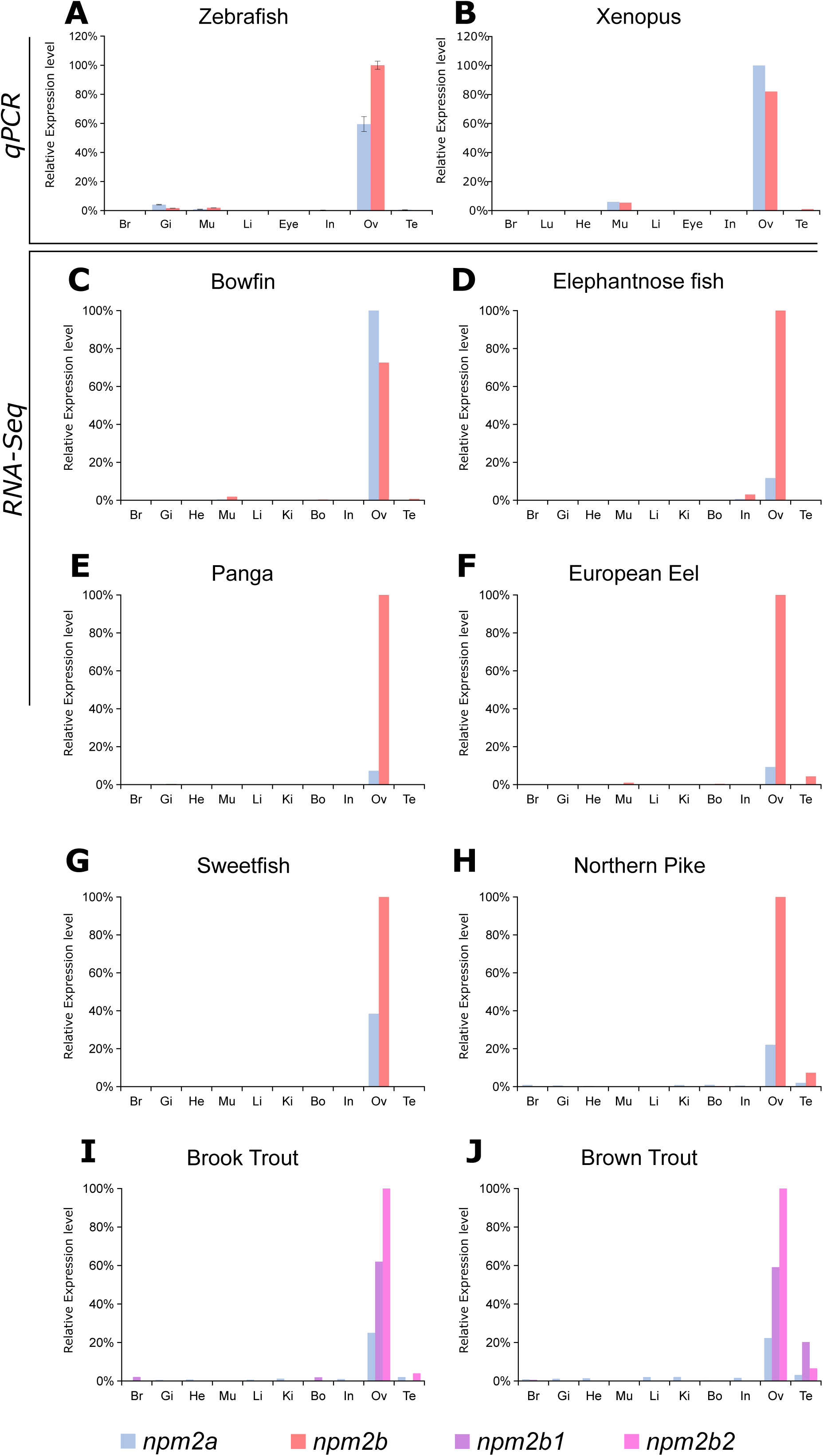
Tissue distribution of *npm2a* and *npm2b* in different species. **A)** Tissue expression analysis by QPCR of *npm2a* and *npm2b* mRNAs in zebrafish and Xenopus. **(B)** Expression level is expressed as a percentage of the expression in the ovary for the most expressed gene. Data were normalized using *18S* expression. **(C-J)** Tissue expression level by RNA-Seq of *npm2a* and *npm2b* mRNAs in different fish species. mRNA levels are expressed in read per kilobase per million reads (RPKM). In salmonids (**I** and **J**), the two 4R isoforms of *npm2b* are *npm2b1* and *npm2b2*. Br, brain ; Gi, gills ; Lu, lung ; In, intestine ; Li, liver ; Mu, muscle ; He, heart ; Bo, bone ; Ki, kidney ; Ov, ovary ; and Te, testis.

### Embryonic expression of npm2a and npm2b

Thus, in order to delve deeper into their functions in reproduction and embryogenesis, we investigated *npm2a* and *npm2b* mRNA expression during oogenesis and early development in zebrafish. During zebrafish oogenesis, both *npm2a* and *npm2b* transcripts were found at high levels in oocytes (Fig. 5A), and despite gradual decreases in the levels of both *npm2* during oogenesis, they still can be detected at reasonable amounts in the unfertilized egg (i.e. metaphase 2 oocyte) (Fig. 5B). Thereafter, *npm2a* and *npm2b* transcript levels progressively decreased during embryonic development after fertilization before reaching very low levels at 24 hours post-fertilization (hpf) (Fig. 5B). This suggests that both mRNAs are strictly maternal (*i.e.* not re-expressed by the zygote), which is consistent with previous studies on zebrafish *npm2a* and *npm2b* transcripts [8,29]. Their expression profiles are typical features of maternally-inherited mRNAs, which highly suggested that the novel *npm2a* is also a maternal-effect gene.

**Figure 5:**
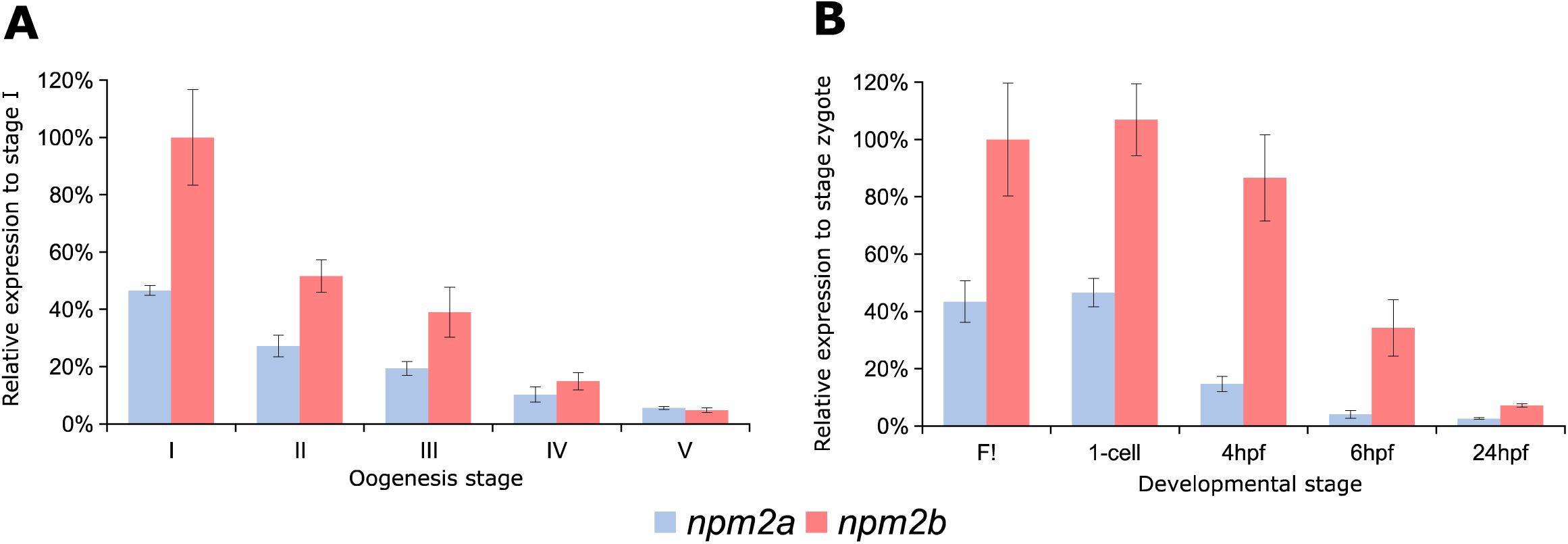
*npm2a* and *npm2b* expression during oogenesis and early development. QPCR analysis of **(A)** *npm2a* and **(B)** *npm2b* expression during oogenesis and early embryonic development in zebrafish. Data were normalized using luciferase, and relative expression was based on *npm2b* expression at the indicated stage. UF, unfertilized egg; hpf, hours post-fertilization.

### Peptidic domains of npm2 paralogs

We further examined the functions of the Npm2 proteins by analysis of their protein domains. To date, only the role of *npm2b* has been investigated in *Xenopus tropicalis* [12,30,31], mouse [14,32], humans [33–35], cattle [27], and zebrafish [8]. As previously demonstrated, *npm2b* is a maternal-effect gene whose transcripts are accumulated in the growing oocyte and maternally-inherited by the zygote, where it functions as a histone chaperone to decondense sperm DNA as well as reorganize chromatin, thus, it has also been suggested to contribute to ZGA as well [8,14,27,28]. Npm2b is thought to be activated by various post-translational modifications and homo-pentamerisation [36], and subsequently interacts with chromatin by exchanging sperm-specific basic proteins with histones via its core domain and acidic tracts A1, A2 and A3 [34,37–42]. However, no data is available on the structure and function of Npm2a. Using a predictive approach, we identified the presence of the Npm core domain, acidic tract A1, and acidic tract A2 in the various investigated Npm2 sequences. The nucleoplasmin family is defined by the presence of an Npm core domain, which enables oligomerization of Npm proteins, in all of its members [38,43]. We were able to predict the presence of this domain in all investigated Npm2a and Npm2b sequences (Supplemental Fig. S1), suggesting that Npm2a proteins could also form homo- or hetero-polymers. The acidic tract domains were demonstrated to facilitate histone binding by increasing the recognition and affinity for different histones [40,44]. Acidic tract A1 was demonstrated to be absent from most of the Npm2b investigated so far, except *Xenopus tropicalis* Npm2b, and acidic tract A2 was predicted to be in all investigated Npm2b apart from guinea pig Npm2b and American whitefish Npm2b1 (Supplemental Fig. S1). This strongly suggested that the histone and basic protein binding activity of Npm2b is mediated predominantly by acidic tract A2. On the other hand, in all investigated Npm2a proteins, we predicted an acidic tract A1 except in that of dogfish, zebrafish, and allis shad (Supplemental Fig. S1). In contrast, only half of the investigated Npm2a proteins harbored an acidic tract A2, which is additionally shorter than the one present in Npm2b proteins (Supplemental Fig. S1). Our results demonstrated that all Npm2a proteins, except those of Allis shad and dogfish, possess acidic tracts, which could potentially mediate histone and basic protein interactions.

### Functional analysis of npm2a and npm2b in zebrafish

Lastly, we performed functional analysis of these two Npm2 proteins by genetic knockout using the CRISPR/cas9 system. One-cell staged embryos were injected with the CRISPR/cas9 guides that targeted either *npm2a* or *npm2b* and allowed to grow to adulthood. Mosaic founder mutant females (F0) were identified by fin clip genotyping and subsequently mated with wild-type (WT) males, and embryonic development of the F1 fertilized eggs was recorded. Since the mutagenesis efficiency of the CRISPR/cas9 system was very high, as previously described [45,46], the *npm2* genes were sufficiently knocked-out even in the transgenic mosaic F0 females. This was evidenced by the substantially lower transcript levels of *npm2a* and *npm2b* in the F1 embryos as compared to those from control WT pairings (Fig.6A). Thus, the phenotypes of *npm2a* (n=3) and *npm2b* (n=4) mutants could be observed even in the F0 generation. Since none of the mutated genes were transmissible to future generations neither through the male nor the female, therefore, all of our observations were obtained from the F0 generation. We observed that most of the embryos from the *npm2b* mutant females underwent cellular division during the very early stages of development (1-3 hpf) despite a considerable number of embryos with abnormal morphology (24.00±7.84% versus 0% in controls) (Fig.6B), which included smaller size, enlarged yolk to membrane ratio, and small yolk to membrane ratio. The diameter of the embryo is demonstrated by the red dotted lines at oblong and germ ring stages, which reveal the extremely reduced size of *npm2a* embryos (Fig.6C). Notably, around two thirds of the embryos showed abnormal cell division, even in those with normal morphology, that culminated in developmental arrest at around 4 hpf following which the cells stopped dividing and appeared to regress. Thus, the developmental success at 4 hpf (oblong/sphere stage) was 38.09±7.31% vs. 85.09±4.88% in controls (Fig.6B and 6C [right column]). By the 24 somite stage (24 hpf), these embryos were all dead while the remaining embryos that showed normal development and normal cell division continued to progress normally. This finding corresponded to our previous results, which showed that *npm2b*-deficient embryos targeted by morpholino arrested at 4 hpf and eventually died [8].

**Figure 6:**
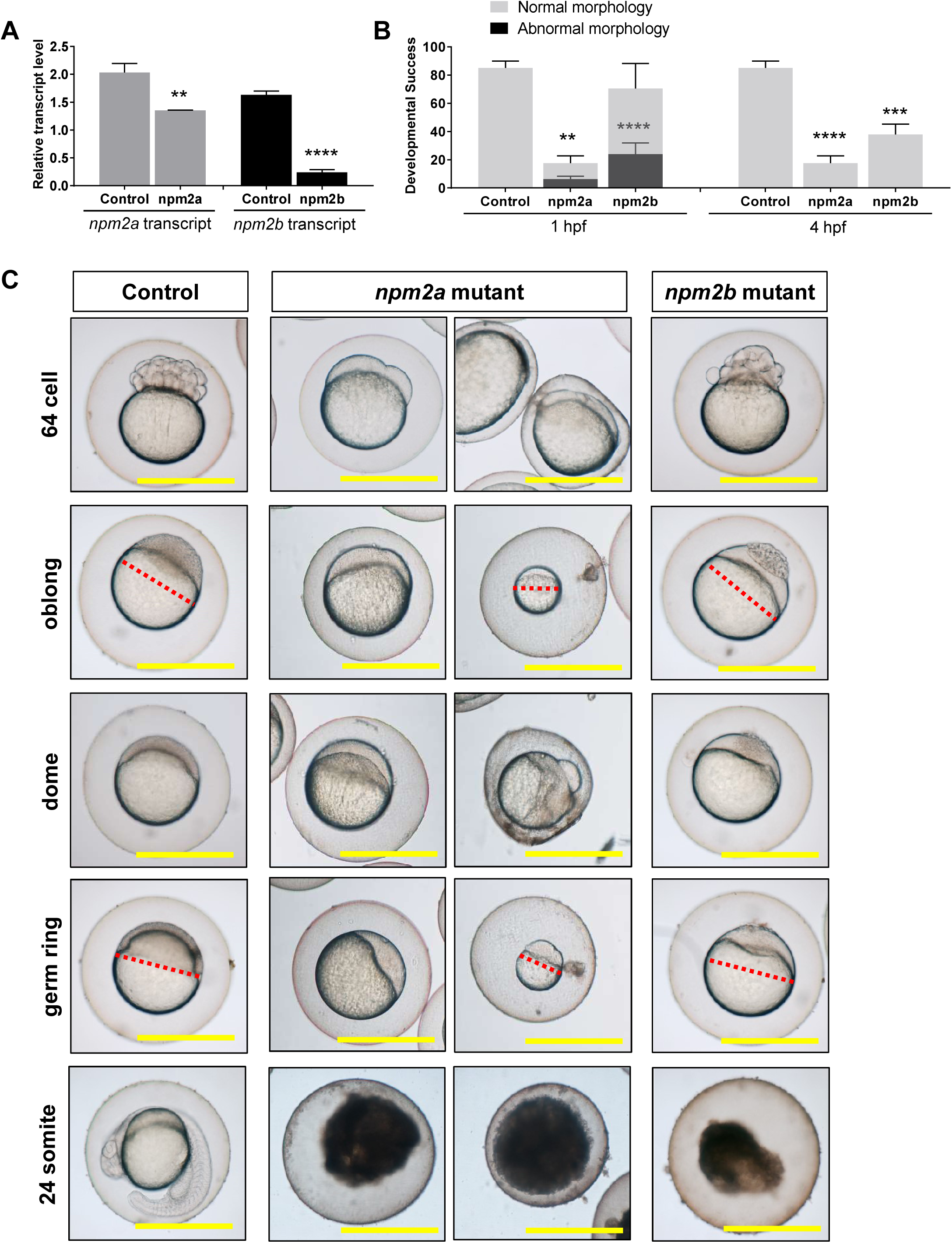
CRISPR/cas9 knockout of *npm2a* and *npm2b* in zebrafish. **(A)** Relative expression level of *npm2a* and *npm2b* transcripts by QPCR in the fertilized zebrafish eggs from crosses between *npm2a* or *npm2b* mutant female and wildtype (WT) male, respectively. **(B)** Developmental success as measured by the proportion of fertilized eggs that underwent normal cell division and reached normal developmental milestones based on Kimmel et al. [51] from crosses between WT animals (control), and *npm2a* or *npm2b* mutant female and WT male at 1 and 4 hours post-fertilization (hpf). Shaded grey columns denote normal morphology and black columns denote abnormal morphology of the fertilized eggs at 1 hpf. **(C)** Representative images demonstrating development of fertilized eggs from crosses between control animals (left panels), and *npm2a* (middle panels) or *npm2b* (right panels) mutant female and WT male at 64-cell, oblong, dome, germ ring, and 24-somite stages according to Kimmel et al [51]. N=3 for *npm2a* mutant and N=4 for *npm2b* mutant. All assessments were performed from at least 3 clutches from each mutant. Scale bars denote 500 nm. Red dotted lines define the diameter of the embryo. ** p<0.01, ***p<0.001, ****p<<0.0001

On the other hand, *npm2a* mutant-derived embryos had a very low developmental success even at a very early stage of growth (1 hpf) (17.62±5.06% vs. 85.09±4.88% in controls) as defined by a complete lack of cell division, and a large population of them (36.36±6.98% vs. 0% in controls) had an abnormal morphology (Fig.6B and 6C [middle columns]). The abnormal morphology of these embryos was similar to that found in *npm2b-*deficient embryos. The F1 embryos that did not undergo any cell division at 1 hpf continued to display a complete lack of development at 64-cell, oblong, dome, germ ring, and finally somite stages (Fig.6B [middle columns]). Similar to the *npm2b* mutant-derived embryos, the *npm2a*-deficient embryos were all dead by 24 hpf while the remaining embryos that showed normal development and normal cell division continued to progress normally. This novel finding showed for the first time that *npm2a* is essential for early development of embryos, and is therefore a crucial maternal-effect gene. Further, we demonstrated that while *npm2a* and *npm2b* share similar tissue distribution, as both are found specifically in the ovaries and early stage embryos, their roles are distinct and essential to embryogenesis, and one could not compensate for the other. Notably, *npm2a* probably plays a major role at or immediately after fertilization since the embryos are arrested before the first mitotic division, which is in line with a role in sperm DNA decondensation since lack of proper processing of the paternal-derived DNA would lead to developmental arrest at the earliest stage and embryonic death. In contrast, *npm2b* appears to function at a later stage or other factors can replace its role in the early stages as the *npm2b*-deficient embryos are capable of dividing until around 4 hpf, which corresponds to previous reports [8,14]. These findings are in line with those demonstrated for other maternal-effect genes in mammals, such as *mater*, *floped*, *filia*, *tle6* [47], and *bcas2* [48] as well as in zebrafish, including *fue* [49] and *cellular atoll* [50], all of which function to moderate the early events of fertilization and embryogenesis and their ablation leads to arrest and mortality at the earliest stages of embryogenesis.

In consequence, the dominancy of the mutant allele combined with the strong maternal effect of these genes, which led to massive early embryonic mortality such that none of the mutant carriers survived, very likely contributed to the lack of germline transmission of the mutations despite reported heritability rates of 10-30% in CRISPR/cas9-targeted zebrafish from previous studies [45,46]. This was supported by detection of the mutant allele via PCR genotyping in the genome of F1 *npm2a*- and *npm2b*-deficient embryos that were developmentally-arrested at the germ ring stage, but not in the surviving animals at 24 hpf (Supplemental Fig.S2).

## Conclusions

Since two annotated genes corresponding to *npm2*, which is known as an essential maternal-effect gene in mammals and amphibians, were recently found in zebrafish, we set out to investigate their evolution and function. In this study, we demonstrated that the two duplicates of *npm2*, *npm2a* and *npm2b*, exist in a wide range of vertebrate species, including ray-finned fish, amphibians, and bird. We also found that the mammalian *npm2* gene is in fact an ortholog of *npm2b*. Using phylogeny and synteny analyses, we traced the origins of those two duplicates to the early stages of vertebrate evolution. Our findings indicate that *npm2a* and *npm2b* genes resulted from a local gene duplication that may have occurred between VGD2 and the divergence of ray-finned fish and tetrapods (~450-500 Mya). This ancient origin is in line by the low sequence identity between the two *npm2* genes, although the main protein domains remain conserved. Both genes exhibit a strict ovarian expression and corresponding transcripts are maternally-inherited in fish and amphibians. Moreover, we demonstrated and confirmed by CRISPR/cas9-directed genetic knockout of *npm2a* and *npm2b* that they are both maternal-effect genes that play essential, but distinct, roles in early embryogenesis. *npm2a* probably plays a major role at or immediately after fertilization, and most likely in chromatin sperm decondensation. In contrast, *npm2b* appears to function at a later stage and could participate in ZGA. Our findings will help us gain further insight into the evolutionary diversity of maternal-effect genes and understand the underlying mechanisms that contribute to reproductive success in vertebrates.

## Material and Methods

### Genomic databases

The following genomic data were extracted and investigated from the ENSEMBL genomic database (http://www.ensembl.org/index.html): human, *Homo sapiens*; mouse, *Mus musculus*; chicken, *Gallus gallus*; Xenopus, *Xenopus tropicalis*; coelacanth, *Latimeria chalumnae*; spotted gar, *Lepisosteus oculatus*; zebrafish, *Danio rerio*; and tetraodon, *Tetraodon nigroviridis*. The Chinese alligator (*Alligator sinensis*) genome was extracted and investigated from the NCBI genomic database (http://www.ncbi.nlm.nih.gov/genome/22419). The *Xenopus laevis* L and S genomes were analyzed from the Xenbase database (www.xenbase.org).

### Transcriptomic databases

The following actinopterygian transcriptomes were retrieved and investigated from the Phylofish database (http://phylofish.sigenae.org/index.html): bowfin, *Amia calva*; spotted gar, *Lepisosteus oculatus*; elephantnose fish, *Gnathonemus petersi*; arowana, *Osteoglossum bicirrhosum*; butterflyfish, *Pantodon buchholzi*; European eel, *Anguilla anguilla*; rainbow trout, *Oncorhynchus mykiss*; allis shad, *Alosa alosa*; zebrafish, *Danio rerio*; panga, *Pangasius hypophthalmus*; northern pike, *Esox lucius*; grayling, *Thymallus thymallus*; Atlantic cod, *Gadhus morua*; medaka, *Oryzias latipes*; European perch, *Perca fluviatilis*; brown trout, *Salmo trutta*; European whitefish, *Coregonus lavaretus*; brook trout, *Salvelinus fontinalis*; Astyanax, *Astyanax mexicanus*; lake whitefish, *Coregonus clupeaformis*; eastern mudminnow, *Umbra pygmae*, and sweetfish, *Plecoglossus altivelis*.

### Gene predictions

#### TBLASTN search

Genomic data were analyzed using the TBLASTN algorithm (search sensitivity: near exact match short) on the ENSEMBL website or the NCBI browser for the Chinese alligator genome. The TBLASTN algorithm on the SIGENAE platform was used on the transcriptomic data.

#### Predictions of *npm2* genes

The peptidic sequences of zebrafish Npm2a and Npm2b were used as query in TBLASTN search to identify the open reading frame (ORF) encoding *npm2* genes in the various investigated genomes and transcriptomes.

### Phylogenetic analysis

Amino acid sequences of 75 Npm2, 3 Npm3, and 3 Npm1 proteins were first aligned using ClustalW. The JTT (Jones, Taylor, and Thornton) protein substitution matrix of the resulting alignments was determined using ProTest software. Phylogenetic analysis of Npm proteins was performed using the Maximum Likelihood method (MEGA 5.1 software) with 1,000 bootstrap replicates.

### Synteny analyses

Synteny maps of the conserved genomic regions in human, mouse, chicken, Xenopus, coelacanth, spotted gar, zebrafish, and tetraodon were produced using PhyloView on the Genomicus v75.01 website (http://www.genomicus.biologie.ens.fr/genomicus-75.01/cgi-bin/search.pl). Synteny analysis of the Chinese alligator conserved genomic regions was performed using TBLASTN searches in the corresponding genomic database. For each gene, the peptidic sequences of human and chicken were used as query, as far as they were referenced in the databases.

### RNA-seq

RNA-seq data were deposited into Sequence Read Archive (SRA) of NCBI under accession references SRP044781-84, SRP045138, SRP045098-103, and SRP045140-146. The construction of sequencing libraries, data capture and processing, sequence assembly, mapping, and interpretation of read counts were all performed as previously reported with some modifications [26]. In order to study the transcript expression patterns and levels of *npm2* for each actinopterygian species presenting 2 or 3 *npm2* genes, we mapped the double stranded RNA-seq reads to the corresponding *npm2* CDS using BWA-Bowtie with stringent mapping parameters (maximum number of allowed mismatches −aln 2). Mapped reads were counted using the idxstat command in SAMtools, with a minimum alignment quality value (−q 30) to discard ambiguous mapping reads. For each species, the number of mapped reads was then normalized for each *npm2* gene across the 11 tissues using RPKM normalization.

### Quantitative real-time PCR (QPCR)

For each sample, total RNA was extracted using Tri-Reagent (Molecular Research Center, Cincinnati, OH) according to the manufacturer’s instructions. Reverse transcription (RT) was performed using 1 μg of RNA from each sample with M-MLV reverse transcriptase and random hexamers (Promega, Madison, WI). Briefly, RNA and dNTP were denatured for 6 min at 70°C and then chilled on ice for 5 min before addition of the RT reagents. RT was performed at 37°C for 1 h and 15 min followed by a 15-min incubation step at 70°C. Control reactions were run without reverse transcriptase and used as negative control in the QPCR study. For each studied tissue, cDNA originating from three individual fish were pooled and subsequently used for QPCR. Follicular oocytes at different stages of oogenesis were obtained from at least 3 different wildtype animals, and pooled embryos were obtained from at least 3 clutches from each individual mutant. QPCR experiments were performed with the Fast-SYBR GREEN fluorophore kit (Applied Biosystems, Foster City, CA) as per the manufacturer’s instructions using 200 nM of each primer in order to keep PCR efficiency between 90% and 100%, and an Applied Biosystems StepOne Plus instrument. RT products, including control reactions, were diluted 1/25, and 4 μl of each sample were used for each PCR. All QPCR were performed in triplicate. The relative abundance of target cDNA was calculated from a standard curve of serially diluted pooled cDNA and normalized to 18S, β-actin, β-2 microglobulin, and EF1α transcripts. The primer sequences can be found in Supplemental Table S5.

### Comparison of npm2a and npm2b peptidic sequences

For the species harbouring Npm2a and Npm2b, or Npm2a, Npm2b1, and Npm2b2, we compared the corresponding protein sequences using EMBOSS Matcher tool on the EBI website (http://www.ebi.ac.uk/Tools/psa/emboss_matcher/). Results of the pair-wise comparisons are presented as identity percentages in Supplemental Table S2. We also estimated the synonymous substitution rates (dS) and non-synonymous substitution rates (dN) between the paralogous Npm2a and Npm2b coding sequences (CDS) using JCoDA v1.14 software (http://www.tcnj.edu/~nayaklab/jcoda). The alignment options were set to “clustal”, and the Yang and Nielsen dN/dS substitution models were used.

### CrispR-cas9 genetic knockout

CRISPR/cas9 guide RNA (gRNA) were designed using the ZiFiT online software and were made against 3 targets within each gene to generate large genomic deletions, ranging from 250-1600 base pairs, that span exons which allow the formation of non-functional proteins. Nucleotide sequences containing the gRNA were ordered, annealed together, and cloned into the DR274 plasmid. *In vitro* transcription of the gRNA from the T7 initiation site was performed using the Maxiscript T7 kit (Applied Biosystems), and their purity and integrity were assessed using the Agilent RNA 6000 Nano Assay kit and 2100 Bioanalyzer (Agilent Technologies, Santa Clara, CA). Zebrafish embryos at the one-cell stage were micro-injected with approximately 30-40 pg of each CRISPR/cas9 guide along with 8-9 nM of purified cas9 protein (a generous gift from Dr. Anne de Cian from the National Museum of Natural History in Paris, France). The embryos were allowed to grow to adulthood, and genotyped using fin clip and PCR that detected the deleted regions. The PCR bands of the mutants were then sent for sequencing to verify the deletion. Once confirmed, the mutant females were mated with wildtype males to produce F1 embryos, whose phenotypes were subsequently recorded. Images were captured with a Carl Zeiss microscope (Jena, Germany) and ToupCam camera (ToupTek, Hangzhou, China).

### Genotyping by PCR

Fin clips were harvested from animals under anesthesia (0.1% phenoxyethanol) and lysed with 5% chelex containing 100 μg of proteinase K at 55°C for 2 hrs and then 99°C for 10 minutes. The extracted DNA was subjected to PCR using Advantage2 system (Clontech, Mountain View, CA) for *npm2b* and Jumpstart Taq polymerase (Sigma-Aldrich, St. Louis, MO) for *npm2a*. The primers are listed in Supplemental Table S5.

### Statistical Analysis

Comparison of two groups was performed using the GraphPad Prism statistical software (La Jolla, CA), and either the Student’s t-test or Mann-Whitney U-test was conducted depending on the normality of the groups based on the Anderson-Darling test.

## Acknowledgements

This work was supported by ANR grants PHYLOFISH (ANR-10-GENM-017) and Maternal Legacy (ANR-13-BSV7-0015) to JB. Authors would like to thank Dr. Yann Audic for providing Xenopus tropicalis animals and Ms. Amélie Patinote for zebrafish rearing and egg production.

## Supplementary Data

**Supplemental Table S1:** list of Npm2 protein sequences used in the phylogenetic analysis.

**Supplemental Table S2:** Npm2a and Npm2b protein sequence comparisons.

**Supplemental Table S3:** list of the genes from the conserved syntenic region of *npm2* genes.

**Supplemental Table S4:** Estimation of the ratio of substitution rates (dN/dS) in *npm2a* and *npm2b* genes.

**Supplemental Table S5:** QPCR and PCR primers.

**Supplemental Figure S1:**
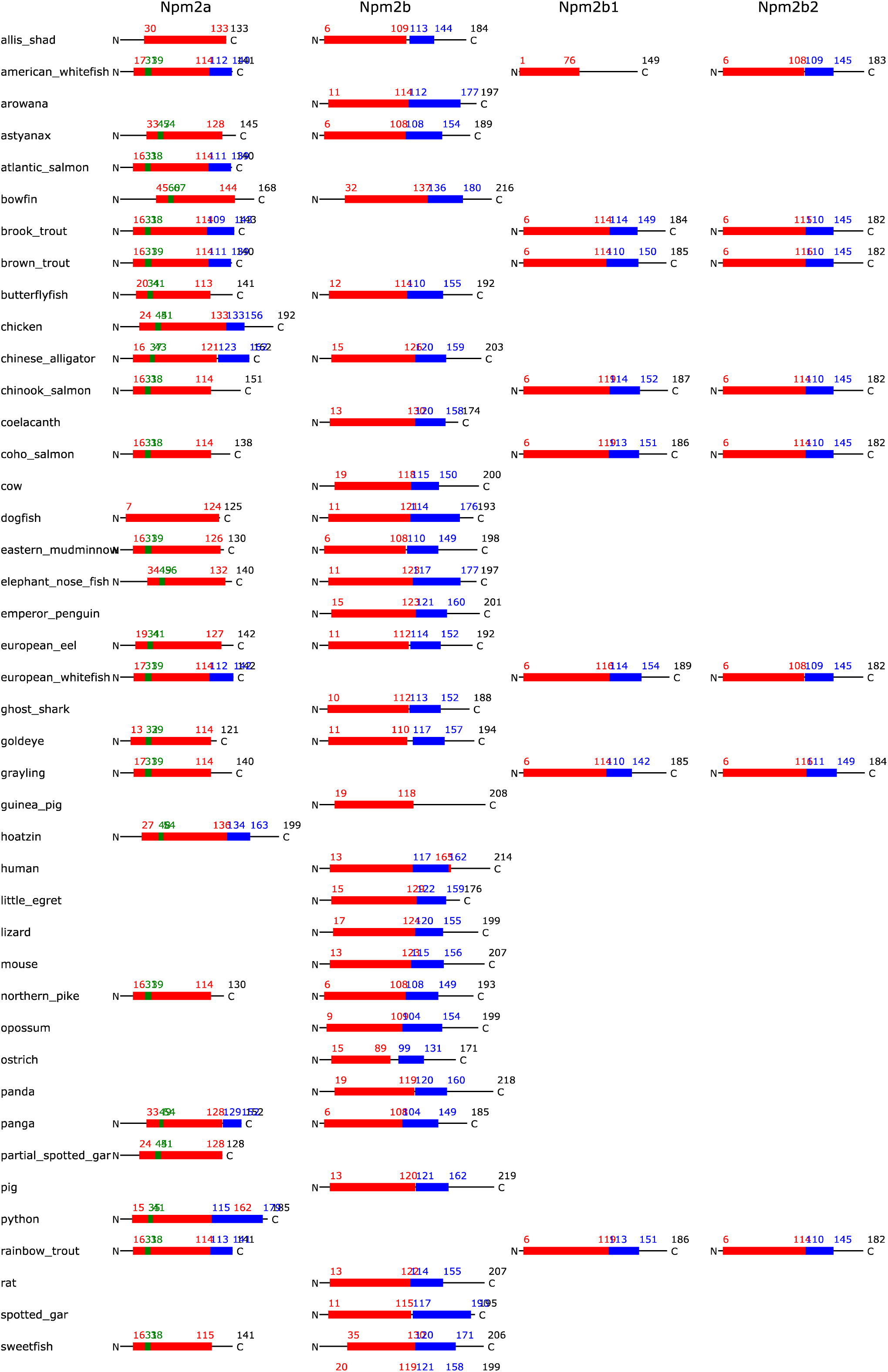
Npm2a and Npm2b protein domain conservation. Presence of the Npm core (red color bar) and acidic tract (blue color bar) domains in *npm2a* and *npm2b* types in various vertebrate species.

**Supplemental Figure S2:**
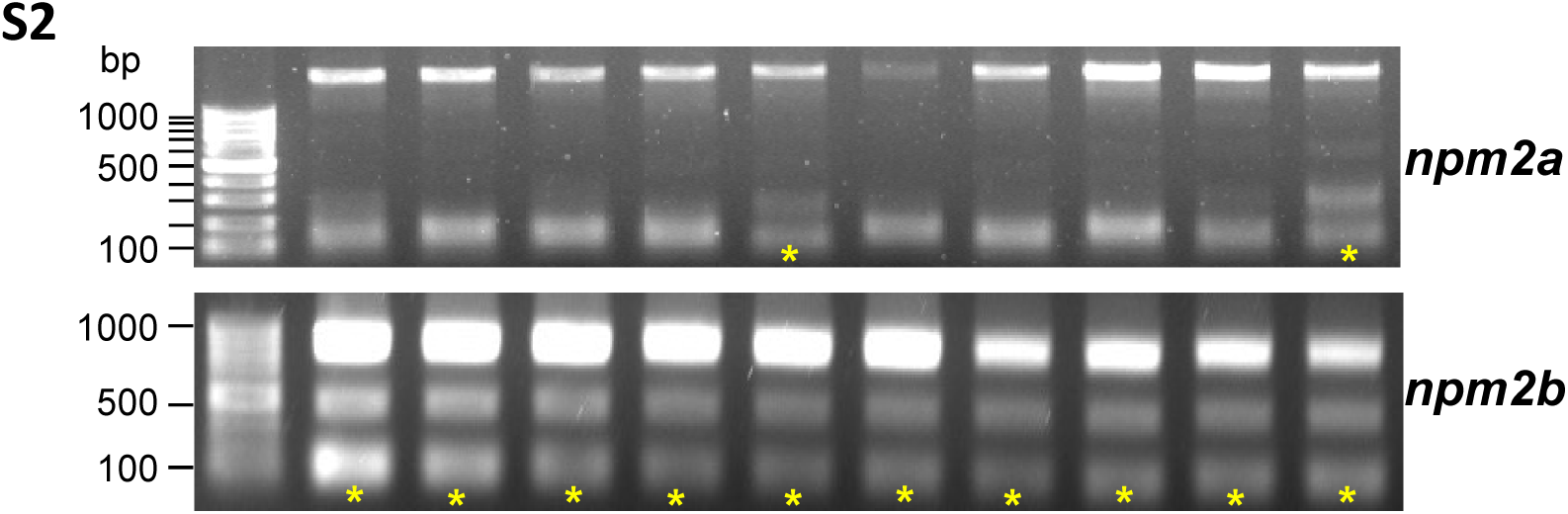
Genotyping by PCR for *npm2a* and *npm2b* in F1 embryos. *npm2a* and *npm2b* knockout animals were crossed with wildtype (WT) zebrafish, and individual F1 embryos were harvested at 6 hours post-fertilization and subjected to genotyping by PCR with gene-specific primers. The wildtype PCR product for *npm2a* was 1634 base pairs (bp) and the CRISPR/cas9-targeted knockout mutant band was 240 bp while the WT *npm2b* band was 813 bp and the CRISPR/cas9-targeted knockout mutant band was 560 bp. The yellow asterisks denote the embryos carrying the mutant allele.

